# A comparative analysis of urinary microbiome identifies putative probiotics

**DOI:** 10.64898/2026.05.15.725591

**Authors:** Riya Anand, Rishav Sahil, Rakesh Pandey, Pradyot Prakash, Hari Sharan Misra, Ganesh Kumar Maurya

## Abstract

Urinary tract infections (UTIs) are the most prevalent bacterial infections globally, and their management increasingly challenged by antimicrobial resistance (AMR). Probiotics offer a promising approach to mitigate AMR by competitively excluding uropathogens and enhancing host immunity by producing immune modulators. Despite being potential, key gaps persist between the discovery of uroprotective probiotic strains and optimization of formulations for urinary tract delivery. Here, we analyzed the urinary microbiome of UTI patients and healthy individuals to identify potential probiotic candidates for the prevention and management of UTIs. Publicly available 16S rRNA amplicon sequencing data of the urinary tract were processed using a standardized pipeline for sequence quality assessment, taxonomic assignment, and microbial function prediction. Comparative analysis showed a significant shift in microbial composition between UTI patients and healthy controls. The dominated phyla identified included Acidobacteriota, Actinobacteriota, Bacteroidota, Campylobacterota, Cyanobacteria, Firmicutes, Fusobacteriota, Patescibacteria, Proteobacteria, and Synergistota. Overall differential abundance analysis revealed *Escherichia coli* as the predominant UTI-associated species, while *Lactobacillus crispatus* was enriched in healthy samples. Additionally, predictive functional analysis indicated that metabolic pathways associated with beneficial microbes were enriched in the healthy group. Overall, the study highlights the association of distinct urinary microbiome signatures with infection status, which supports *L. crispatus* as the most promising probiotic for UTI prevention and control.

## Introduction

Globally, urinary tract infections (UTIs) present one of the most prevalent bacterial infections, with antimicrobial-resistant (AMR) strains that are frequently encountered in healthcare settings. Healthcare accounts for 40% of all hospital-acquired UTIs. UTIs are classified by severity (uncomplicated or complicated) and by their anatomical location (upper or lower urinary tract) [1]. Infections of the upper urinary tract, including pyelonephritis and ureteritis, tend to be more severe and are generally more serious, often characterized by symptoms such as high-grade fever, flank pain, and urine that is dark or has a strong odour. In contrast, lower UTIs, which involve the bladder (cystitis) and the urethra (urethritis), are generally associated by symptoms like dysuria (painful urination), increased urinary frequency, and urgency [2].

Colonization of the periurethral space by commensal or gut-derived bacteria is the initial step in UTI pathogenesis, followed by bacterial ascent through the urethra and the establishment of infection in the bladder. Upon reaching the bladder, bacteria adhere to the urothelial lining via pili and adhesins, which triggers an inflammatory response. In response to infection, neutrophils migrate into the bladder; however, uropathogens can escape immune detection by forming biofilms. These biofilms serve as a shield, enhancing their persistence within the urinary tract. The secretion of bacterial toxins and proteases compromises the epithelial barrier, facilitating pathogen ascent through the ureters to the kidneys and causing pyelonephritis. If not properly treated, the infection may escalate, leading to bacterial invasion of the bloodstream, known as bacteremia [3]. Overuse of antibiotics, especially for recurrent UTIs, has accelerated the rise of AMR. A major resistance mechanism is the production of extended-spectrum beta-lactamases (ESBLs) that give resistance to cephalosporins and penicillins [4]. Most UTIs are attributed to uropathogenic *E. coli* (UPEC), accounting for 46.4% to 74.2% of isolates, followed by *Klebsiella* spp. (6.0% to 13.45%), *Enterococcus* spp. (5.3% to 9.54%), and *Proteus* spp. (4.7% to 11.9%). UTIs can be distressing and, in severe cases, life-threatening. Approximately 12% of males and 40% of females are reported to develop at least one symptomatic UTI over the course of their lifetime [5].

The Global Burden of Disease Study (2019) reported that UTIs accounted for over 404.6 million incident cases worldwide. The burden of UTIs has increased by 60.4% from 1990 to 2019, with the highest incidence observed in South Asia, Western Europe, and tropical Latin America. Mortality associated with UTIs remains significant, with India recording the highest number of deaths (over 55,000) in 2019. Notably, women are disproportionately affected, with an ASIR (age-standardized incidence rate) 3.6 times higher than men, particularly in the 30-34 year age group. The global rise in antimicrobial resistance, particularly in UPEC, has further exacerbated the health burden and mortality associated with UTIs, underlining the immediate action for better infection control, surveillance, and development of antibiotic alternatives [6,7].

As AMR escalates, conventional antibiotics are becoming less effective. The World Health Organization (WHO) emphasizes a ‘One Health’ strategy to address AMR across human, animal, and environmental health. To reduce antibiotic dependence and resistance, non-antibiotic strategies such as probiotics, plant-derived antimicrobials, and dietary supplements are increasingly being explored. These approaches aim to manage infections while preserving antibiotic efficacy and reducing the clinical and economic burden on healthcare systems [4].

Recent advancements underscore the growing role of probiotics as alternative or adjunct therapies for UTIs, particularly in the context of rising AMR. Clinical trials have established the effectiveness of strains such as *Lactobacillus crispatus* (Lactin-V), *L. rhamnosus* GG and *E. coli* 83972, which have shown promise in clinical trials for UTI prevention [8]. However, limitations persist, including strain-specific variability, poor mucosal persistence, inconsistent clinical outcomes, and a lack of large-scale, standardized randomized controlled trials.

In this study, we performed a comprehensive analysis of the urinary microbiome to explore potential probiotic candidates for UTI management. We analysed amplicon sequencing data from urine samples of UTI patients and healthy individuals sourced from public repositories. Our study involved rigorous quality control, taxonomic classification, and community composition analysis to identify microbial signatures associated with urinary health. Differential abundance analysis identified *E. coli* as significantly enriched in UTI patients, while *L. crispatus* was predominant in healthy individuals. Additionally, microbial-pathway correlation analysis reinforced the role of beneficial taxa in maintaining urinary tract homeostasis. These findings provide insights for the development of probiotic interventions aimed at preventing and managing UTIs.

## Materials and methods

### Data collection

We performed a structured literature search for published microbiome studies related to urinary tract infections (UTIs) using the PubMed (https://pubmed.ncbi.nlm.nih.gov/) and Google Scholar (https://scholar.google.com/) search engines (keywords: “urinary tract infection”, “healthy”, “microbiome”, “metagenome”, “amplicon”, and “16S rRNA”). The search yielded abstracts published between 2011 and 2024.

We screened abstracts and associated metadata to identify studies employing 16S rRNA amplicon sequencing on urine samples from healthy individuals or UTI patients. Studies were included only if they had sequencing data derived from urine and metadata available describing the clinical condition (e.g., infection status). Studies involving animal subjects or lacking sufficient clinical metadata were excluded.

We retrieved datasets from studies via accession numbers. Sample identifiers were linked to metadata files describing clinical status, age, gender, urine collection method, and sequencing parameters. Publicly available databases, such as Sequence Read Archive (SRA) and European Nucleotide Archive (ENA) were used to get the raw data files for 16S amplicon studies, using the following identifiers: PRJNA385350 [9], PRJNA678697 [10], PRJNA686823 [11], PRJNA705267 [12], PRJEB4256 [13], PRJEB2717 [14] and PRJEB20159 [15]. In total, we analyzed 16S rRNA data from 274 urine samples across all study interventions.

### Data processing and taxonomy assignment

The analysis was conducted using a standardized microbiome analysis pipeline implemented by Sahil et al [16]. The pipeline includes quality filtering, taxonomic assignment, and normalization for amplicon sequences, allowing for the comparison of microbial composition and diversity. Firstly, the single-end and paired-end raw sequence data were converted into FASTQ format files. FastQC (v0.12.1) (http://www.bioinformatics.babraham.ac.uk/projects/fastqc/) and MultiQC (v1.22.3) [17] were used for quality assessment of the sequences. The sequencing reads were cleaned by using fastp (v0.23.2) [18]. In the cleaning steps of fastp, reads with poor quality score, adapter contamination, and reads containing ambiguous nucleotides (N) were removed. High-quality sequences were retained for subsequent analyses.

The DADA2 pipeline [19] in R (v4.3.3) was further used for sample inference, filtering, chimera removal, dereplication and merging of paired-end reads. Low-quality sequence bases were manually trimmed from the end of the sequences prior to these steps. Strict dereplication (clustering of duplicate sequences) was applied to the filtered sequences, and Amplicon Sequence Variants (ASVs) were identified and counted.

ASVs were aligned to the SILVA database (v138.1) for taxonomy assignment. Archaeal and mitochondrial sequences were removed from the dataset. ASVs were merged at the species level by computing the mean abundance of all ASVs corresponding to each species. The ASVs that were not annotated at the species level were excluded. The relative abundance was calculated by dividing the abundance of each ASV in a sample by the total abundance of all ASVs in that sample, using the transform function of the microbiome R package [20]. Batch effect adjustment was performed using MMUPHin [21]. Further, downstream analyses were conducted on the resulting species-level relative abundance data.

### Bacterial community composition and diversity

Phylum-level abundance across different studies and conditions was analysed. Stacked barplots at the phylum level for all samples were generated using the ggplot2 R package (v3.5.1) within the phyloseq R package (v1.46.0). The comparison of phylum abundances between conditions was further stratified by gender. Furthermore, differences in relative abundance of bacterial phyla and the Firmicutes-to-Bacteroidota (F/B) ratio between UTI and healthy groups were calculated. Statistical differences in the relative abundance of these genera were assessed using the Wilcoxon rank-sum test. A venn diagram was created to display the UTI and healthy-associated genus and species, as well as those that were found to be shared between both conditions.

Alpha (within-sample) diversity was calculated for all the samples, grouped by studies. Shannon and Simpson indices were used as alpha diversity metrics, implemented in the phyloseq R package [22]. Statistical significance was evaluated with the help of Wilcoxon rank-sum test, and a *p*-value < 0.05 was considered significant.

Beta (between-sample) diversity was assessed using between-classes principal component analysis (PCA) with the R package ade4 (v1.7.22) [23] to evaluate taxonomic differences between healthy and UTI conditions. The statistical significance of between-classes PCA was determined using a Monte Carlo test with 9,999 permutations.

### Pathway prediction

Microbial functional profiles were predicted using non-merged ASV abundance data and representative sequences using PICRUSt2 pipeline [24], based on the MetaCyc database. Predicted pathway abundances from all studies were merged by summing the mean abundance of each pathway.

### Differential abundance analysis

Differential abundance analyses of bacterial species and pathways of two different conditions, UTI and healthy, were conducted using the *edgeR* package (v4.0.16) [25] in R. Linear modeling was performed, followed by contrast fitting to compare UTI against healthy samples. Empirical Bayes moderation was applied to improve variance estimation, and *p*-values were adjusted for multiple testing using the Benjamini-Hochberg method to control the false discovery rate (FDR). Species with adjusted *p*-values (*p* < 0.05) were considered significantly differentially abundant between the two conditions.

### Bacterial communities and pathways correlation analysis

To explore potential functional associations between bacterial species and predicted metabolic pathway abundances, correlation analysis was performed. Spearman’s rank correlation coefficients were computed in the *psych* R package (v2.4.12).

### Additional R packages used

RcolorBrewer v1.1.3, vegan v2.5, factoextra v1.0.7, dplyr v1.1.4, tidyverse v2.0.0, forcats v1.0.0, corrplot v0.95.

## Results

### Bacterial diversity across the multiple samples

We obtained amplicons and corresponding metadata for 162 urine samples from individuals with UTIs and 112 urine samples from healthy individuals, derived from seven publicly available studies. Only human-derived samples were included, and non-human samples were excluded. Different variable regions targeted for 16S rRNA gene sequencing were used among different studies. The compiled metadata (S1 Table) and abundance count data (S2 Table) for all samples are provided in the supplementary material.

We identified 11 different bacterial phyla across all 274 samples and conditions (UTI and healthy), including Actinobacteriota, Bacteroidota, Firmicutes, etc. (Fig 1A and 1B). The phyla showing significant differences between healthy individuals and UTI patients were Acidobacteriota, Bacteroidota, Campylobacterota, Cyanobacteria, Firmicutes, Fusobacteriota, Patescibacteria, Proteobacteria, and Synergistota (Fig 1C). Gender-wise analysis showed similar results (Fig S1A, S1B, S2A and S2B). There was no significant difference in Firmicutes-to-Bacteroidota (F/B). This trend remained consistent when stratified by gender: both female and male UTI samples.

**Fig 1.**
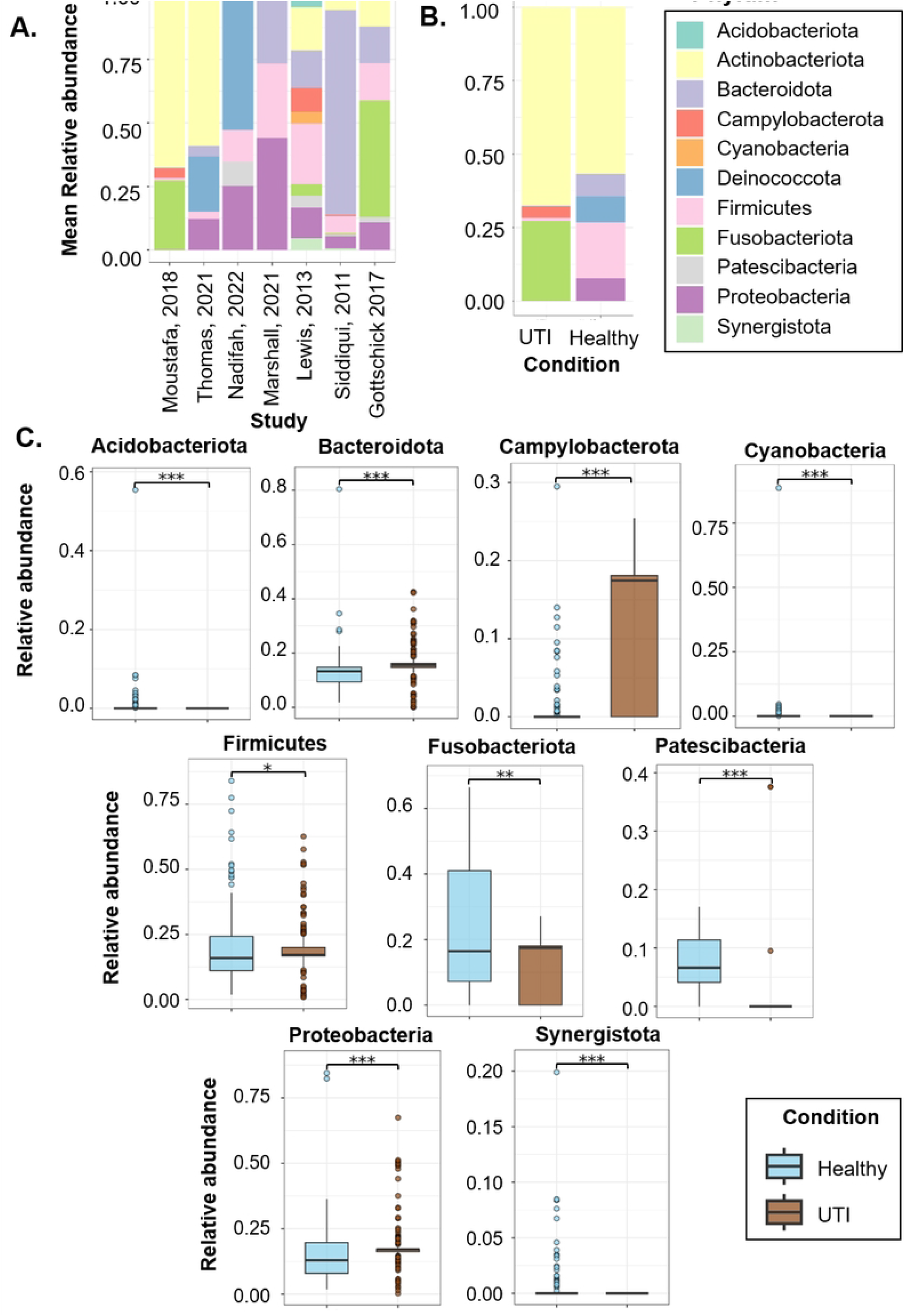
Diversity and composition of the urinary microbiota. (**A**) Stacked bar plot showing phylum-level relative abundance across studies. (**B)** Stacked bar plot comparing phylum-level relative abundance between UTI patients and healthy individuals. (**C)** Boxplot of the relative abundance of significantly abundant phyla in UTI and healthy conditions. Asterisks indicate statistical significance obtained by Wilcoxon rank-sum test (\**p* < 0.05, \*\**p* < 0.001, \*\*\**p* < 0.0001).

### Distinct bacterial identification between UTI and healthy conditions

The bacterial composition across all the samples revealed 113 genera unique to healthy individuals, 24 genera exclusive to UTI cases, and 46 genera common to both groups. Healthy females exhibited 124 unique genera compared to 25 found exclusively in UTI females, with 26 genera common between them. Similarly, 96 genera were unique to healthy males, 24 were specific to males with UTIs, and 46 genera were common to both groups (Fig 2A).

**Fig 2.**
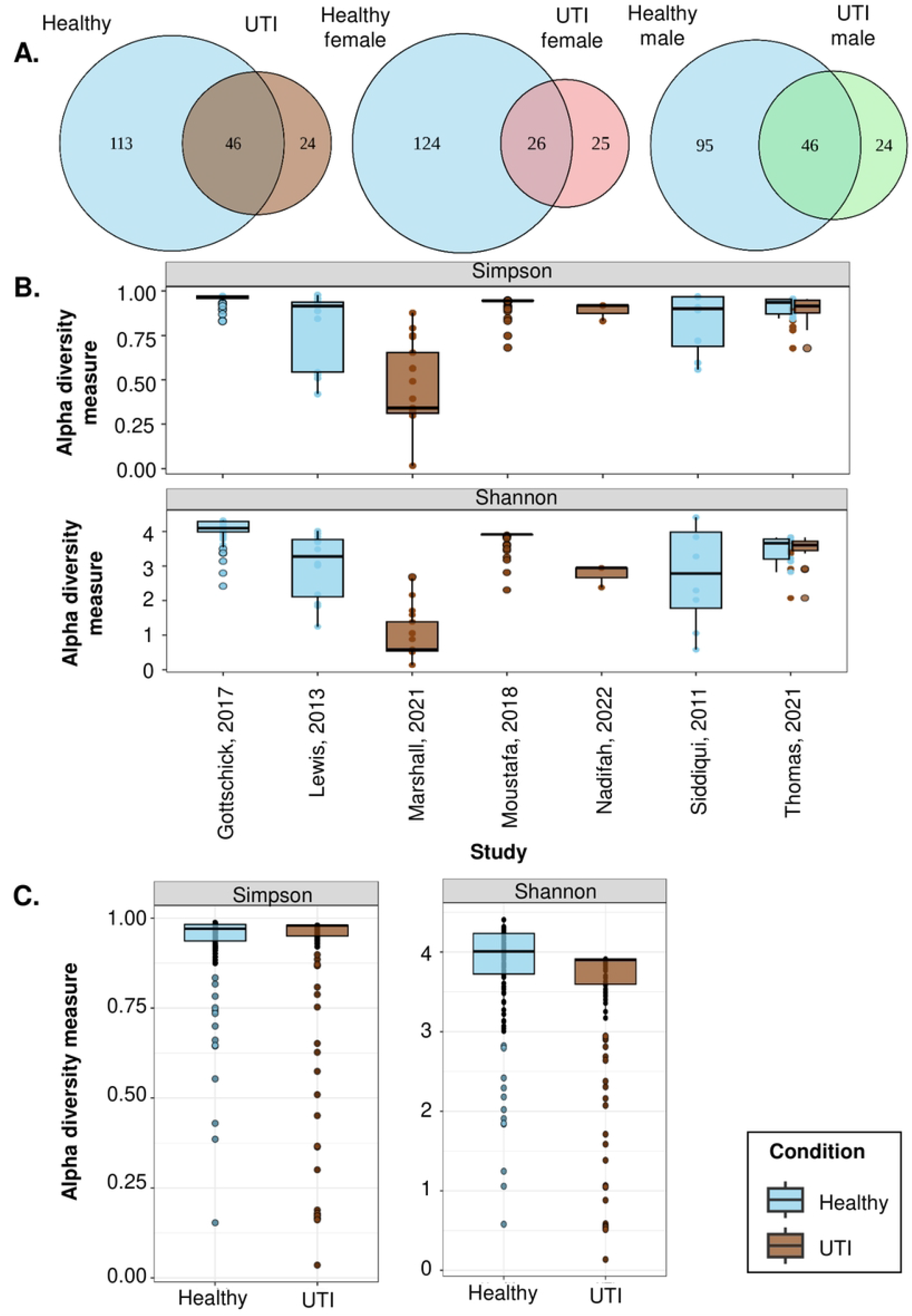
Genus distribution and diversity of the urinary microbiota. **(A)** Venn diagram of genus distribution in healthy and UTI, healthy females and UTI females, and healthy males and UTI males. **(B)** Alpha diversity across different studies. **(C)** Alpha diversity among UTI patients and healthy individuals.

Analysis of species distribution displayed distinct differences in microbial composition between healthy controls and patients with UTIs. Across all samples, 55 species were shared by both healthy and UTI groups, while 128 species were unique to healthy individuals and 33 were exclusive to UTI samples, suggesting a greater microbial richness in healthy conditions. When stratified by gender, healthy females harboured 135 unique species, with 29 common and 33 exclusives to UTI females. Among males, 108 species were unique to the healthy group, 52 were common, and 36 were found only in UTI samples (Fig 3A).

**Fig 3.**
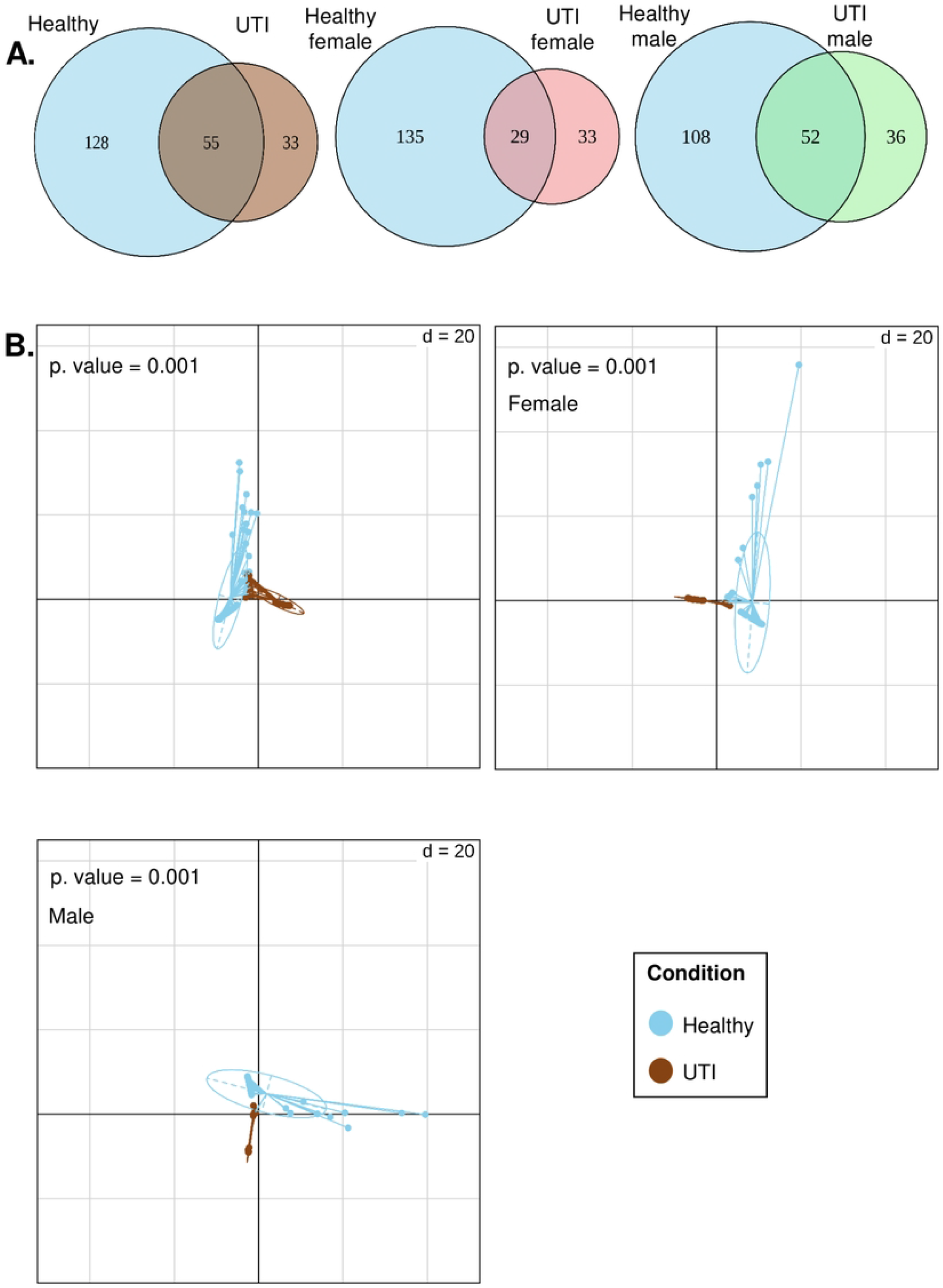
Species distribution and diversity of the urinary microbiota. **(A)** Venn diagram of species distribution in healthy and UTI, healthy females and UTI females, and healthy males and UTI males. (**B)** Beta diversity among healthy and UTI, healthy females and UTI females, and healthy males and UTI males.

Alpha diversity (within-sample diversity) analysis was performed using Shannon and Simpson indices across multiple studies, each with varying sample sizes and study designs (Fig 2B). Only one study, Thomas (2021), included both UTI and healthy control samples, enabling direct within-study comparisons. The remaining studies, Gottschick (2017), Lewis (2013), and Siddiqui (2011) provided data exclusively for healthy individuals, whereas Marshall (2021), Moustafa (2018), and Nadifah (2022) included only UTI samples. This heterogeneity in sample composition and group representation may contribute to observed variability in diversity values across studies. Comparison of Shannon and Simpson indices between healthy individuals and UTI patients showed consistent differences in microbial diversity (Fig 2C). Healthy samples consistently show higher Shannon diversity compared to UTI samples. This suggests that healthy urine microbiomes are more diverse, both in terms of species richness and evenness. The Simpson Index, consistent with the Shannon results, indicates that UTI samples exhibit lower Simpson diversity, suggesting dominance by fewer taxa. In healthy individuals, higher Simpson indices suggest a more balanced microbial community. UTIs are associated with a reduction in microbial diversity and an increase in dominance by specific pathogenic taxa, consistent across studies.

Further exploration of β-diversity in healthy individuals and UTI patients showed microbiome differences, visualized using between-classes PCA. We demonstrated the profiles of community composition for the two groups at the species level, showing significant (*p*-value = 0.001) differences in microbial composition between the two health states (Fig 3B).

### Identification of marker bacteria in UTI and healthy conditions

Differentially abundant bacteria were identified between all UTI patients and healthy controls using the *edgeR* analysis pipeline. 146 differentially abundant species (S3 Table) were found that were significantly different at an adjusted *p*-value < 0.05 with log_2_ fold change (FC). Log_2_ FC < 0 indicates the bacterial taxa are differentially abundant in healthy controls, while log_2_ FC > 0 indicates the bacterial taxa are differentially abundant in UTI samples. We found *E. coli, Gardnerella vaginalis, Porphyromonas somarae, Fusobacterium gonidiaformans*, etc. differentially abundant in UTIs and *L. crispatus, Sneathia amnii, Prevotella amnii, L. iners*, etc. differentially abundant in healthy conditions (Fig 4). Differential analysis in females shows the similar result (Fig S3), but in males, *P. somerae* and *S. amnii* are differentially abundant in UTI and healthy conditions, respectively (Fig S4).

**Fig 4.**
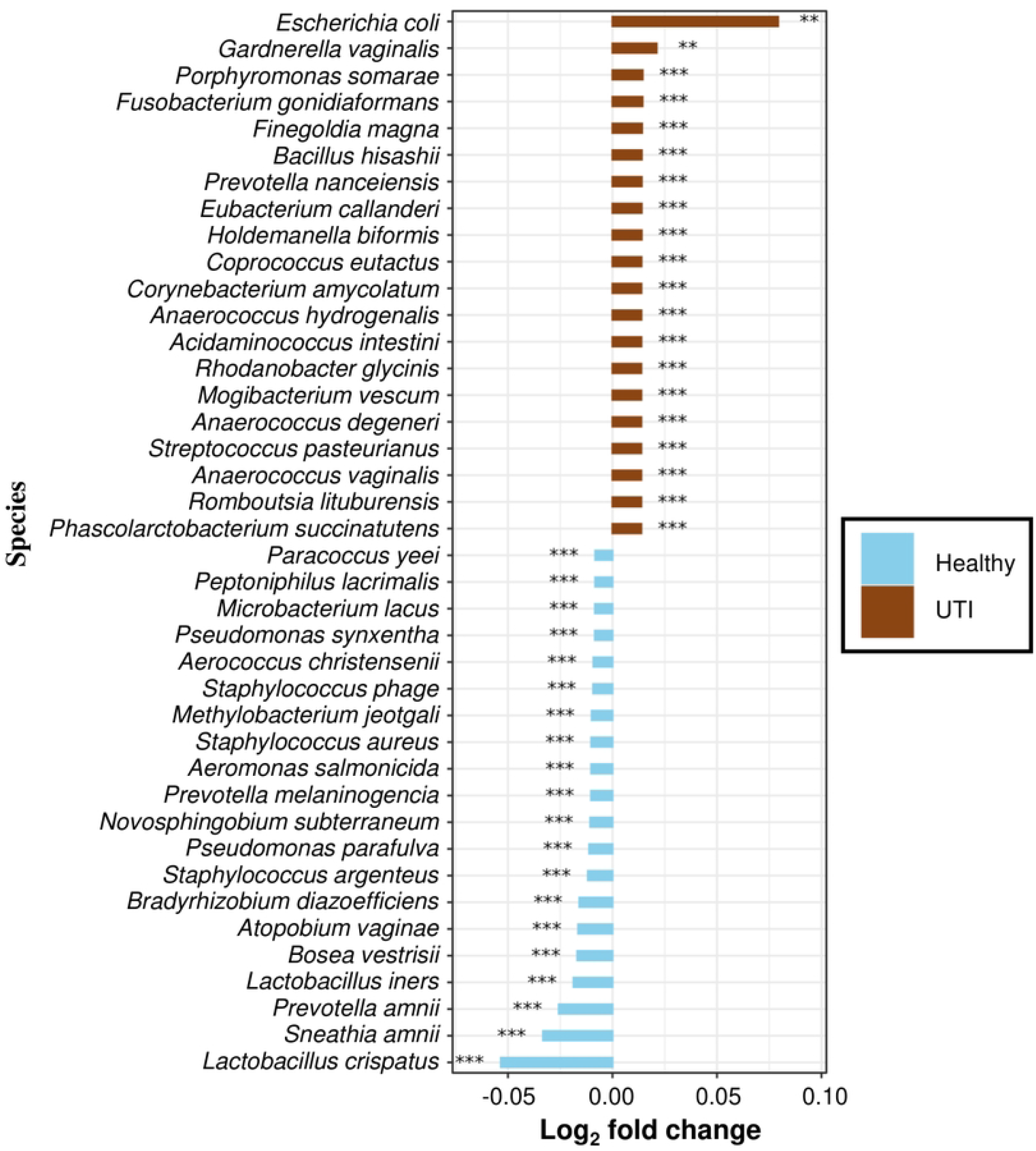
Bar plot of the top differentially abundant microbial taxa in UTI patients and healthy individuals. Log2-transformed relative abundance represents the logarithm of the relative abundance ratio in base 2. Asterisks indicate statistical significance (\**p* < 0.05, \*\**p* < 0.001, \*\*\**p* < 0.0001).

### Functional profiling of marker bacteria

We employed PICRUSt2 for functional profiling of the microbial community using the MetaCyc database. The total pathway we obtained across all studies is 394. Additionally, differentially abundant pathways were identified between all UTI patients and healthy controls, with 304 pathways showing differential abundance (S4 Table). The top 10 differentially abundant pathways of UTIs as well as healthy conditions were annotated (S5 Table).

A Spearman correlation analysis between the most abundant bacterial taxa (top 20 species) and the most represented metabolic pathways, based on 16S rRNA gene amplicon sequencing data from healthy individuals and UTI patients, showed clear differences in microbial functional associations. The correlation plot displayed distinct clusters, with several bacterial species showing strong positive correlations (blue circles) with metabolic pathways in healthy individuals. In contrast, taxa predominant in UTI samples showed predominantly negative correlations (red circles). *E. coli*, despite being the most abundant in UTI conditions, exhibited a correlation pattern more closely resembling that of bacteria enriched in healthy individuals (Fig 5).

**Fig 5.**
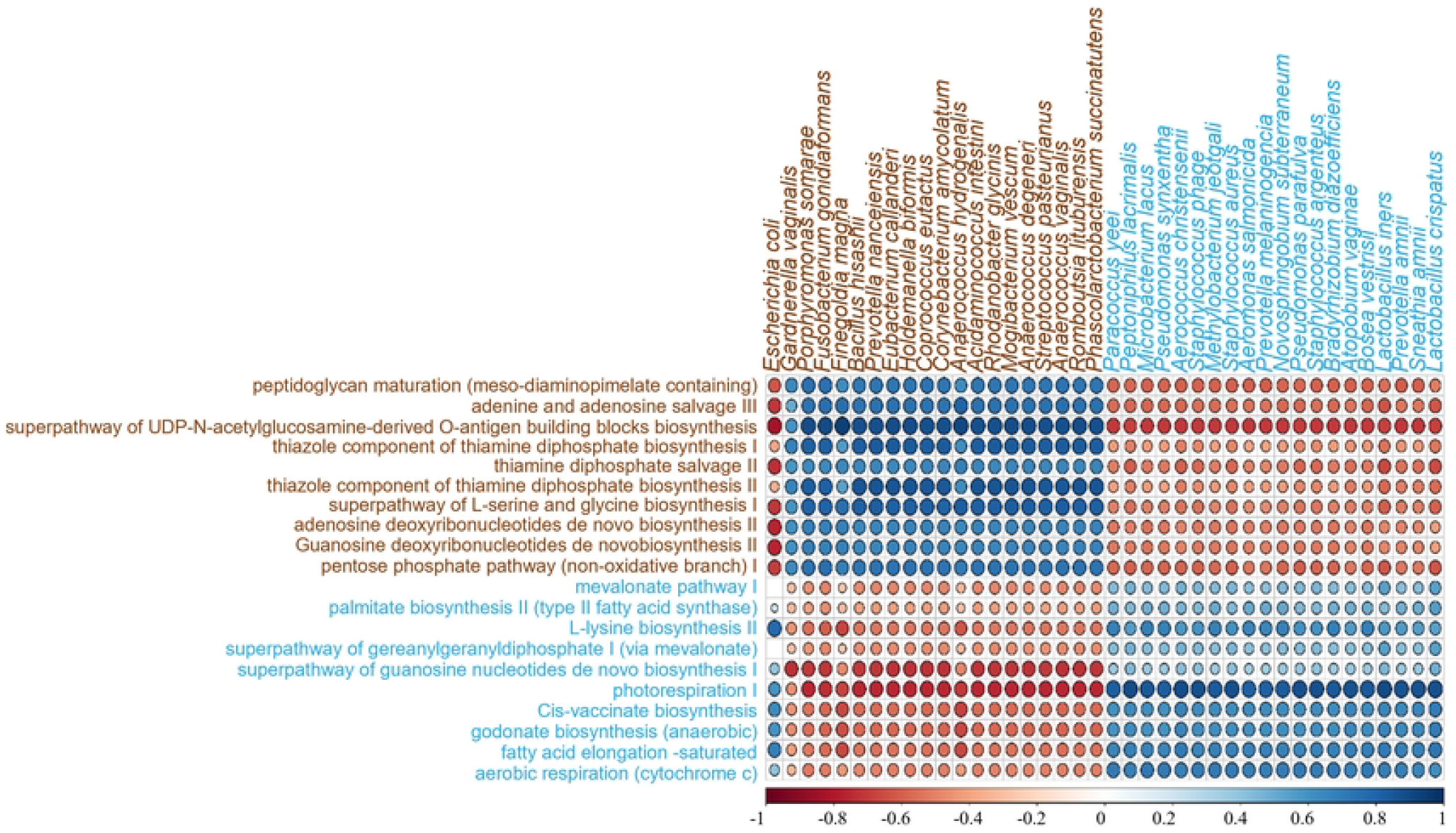
Spearman correlation bubble plot of the top differentially abundant microbial taxa and pathways in UTI patients and healthy individuals.

## Discussion

Many studies comparing individuals with UTIs to healthy controls have demonstrated that the urinary tract microbiota plays a crucial role in maintaining health and is vital in preventing disorders such as UTIs [26–29]. Advancement in DNA sequencing technologies have enabled the identification of bacteria without the need for traditional culture methods. Instead of studying the genome of single bacterial strain grown in a laboratory, the metagenomic approach examines the collective genomes of microbial communities present in natural environments [30]. It is well-documented that a variety of bacteria and fungi can cause UTIs; however, the most common pathogens are uropathogenic *E. coli* (UPEC), *K. pneumoniae, E. faecalis*, and *Proteus mirabilis* [3]. High recurrence rates and rising AMR among uropathogens are likely to significantly increase the economic burden of this infection [3]. In the present study, we analysed publicly available datasets to identify differences in the microbiome of the urinary tract in UTIs and healthy conditions. The datasets included in the current studies are from different geographical locations. Our analysis reveals a significant difference in the microbiome of individuals with UTIs and those who are healthy. In the phylum-level study, we observed a significant change in the phyla, including Acidobacteriota, Bacteroidota, Campylobacterota, Cyanobacteria, Firmicutes, Fusobacteriota, Patescibacteria, Proteobacteria, and Synergistota. Also, we found that there was no significant difference in Firmicutes/Bacteroidetes (F/B) abundance in urine samples from individuals with UTI compared to healthy controls.

Alpha diversity analysis revealed distinct community patterns between UTI and healthy samples. The Shannon index was significantly lower in UTI samples, indicating reduced richness and evenness of the microbial community. This suggests UTI conditions are characterized by overrepresentation of a limited set of genera, consistent with pathogen dominance that reduces overall diversity. In contrast, healthy samples showed higher Shannon values, reflecting a more complex and balanced microbial structure. Interestingly, Simpson index values were slightly higher in UTI samples, implying that although fewer taxa were present, their abundances were more evenly distributed among dominant members. Together, these complementary indices highlight that UTIs are associated with loss of microbial diversity and a shift towards dominance by a narrower set of genera, which may compromise community resilience and facilitate infection. Beta diversity analysis suggests there is a significant difference between the microbiota of UTI patients and healthy controls. The species-level differential abundance analysis demonstrated a clear separation between UTI and healthy samples. In UTIs, the most prominent differentially abundant species are *E. coli*. One of the notable findings in our differential abundance analysis was the enrichment of *L. crispatus* in the healthy group. Apart from *L. crispatus* and *L. iners* all the species differentially abundant in healthy conditions are rare opportunistic pathogens. *L. crispatus* is a well-established beneficial species that plays a key role in maintaining urogenital health through the production of lactate (a major metabolite secreted by lactobacilli), which helps to lower the pH and inhibit the growth of pathogenic organisms [31]. Its protective role has been extensively studied in the vaginal microbiome, where a higher abundance of *L. crispatus* is associated with resistance against urinary tract infections and bacterial vaginosis [32– 34]. Given these attributes, *L. crispatus* has been investigated as a potential probiotic, with strains already explored for intravaginal or oral supplementation to restore microbial balance and reduce infection recurrence.

The Spearman correlation analysis between bacterial taxa and metabolic pathways highlights distinct microbial-functional interactions in healthy individuals compared to UTI patients. The strong positive correlations observed among several taxa and metabolic pathways in healthy samples suggest a functionally stable and cooperative urinary microbiome. These associations likely reflect balanced microbial interactions that contribute to maintaining metabolic homeostasis and host health. In contrast, the predominance of negative correlations in UTI samples indicates a disrupted microbial network, where disease-associated taxa may compete for resources or suppress pathways involved in normal metabolic functioning. Such functional decoupling is a hallmark of microbial dysbiosis commonly observed during infection and inflammation.

Interestingly, *E. coli*, a common uropathogen in UTI patients, displayed correlation patterns resembling those of commensal bacteria in healthy individuals. This finding suggests that not all *E. coli* strains in the urinary tract are pathogenic, supporting emerging evidence of strain-level diversity within the species. Specific *E. coli* lineages may persist as harmless commensals or even contribute to niche stability in the urinary microbiome under non-pathogenic conditions. The observation underscores the importance of considering strain-specific virulence factors, host-microbe interactions, and ecological context when interpreting *E. coli* abundance in urinary microbiome studies.

Overall, these results reinforce the concept that urinary tract health is shaped by both the composition and the functional organization of its microbial community. The shift from positively coordinated metabolic networks in health to negatively correlated, fragmented interactions in disease reflect the metabolic and ecological imbalance associated with UTI. Future studies integrating metagenomic and metabolomic data could further clarify the functional consequences of these correlations and identify key microbial pathways that maintain urinary tract homeostasis or drive infection.

## Acknowledgments

R.A. gratefully acknowledges NBCFDC for NFOBC senior research fellowship. G.K.M. acknowledges IoE, BHU, UGC, India, and SERB-DST, India (SRG/2021/000900) for financial support.

## Supporting information

**S1 Fig. Diversity and composition of the female’s urinary microbiota**. (**A**) Stacked barplot showing phylum-level relative abundance between UTI patients and healthy individuals. (**B**) Boxplot of the relative abundance of significantly abundant phyla in UTI and healthy conditions. Asterisks indicate statistically significant differences by a Wilcoxon rank-sum test (\**p* < 0.05, \*\**p* < 0.001, \*\*\**p* < 0.0001).

**S2 Fig. Diversity and composition of the male’s urinary microbiota**. (**A)** Stacked barplot showing phylum-level relative abundance between UTI patients and healthy individuals. (**B)** Boxplot of the relative abundance of significantly abundant phyla in UTI and healthy conditions. Asterisks indicate statistically significant differences by a Wilcoxon rank-sum test (\**p* < 0.05, \*\**p* < 0.001, \*\*\**p* < 0.0001).

**S3 Fig.** Bar chart showing the top differentially abundant microbial taxa in female UTI patients and female healthy individuals. The log2-transformed relative abundance represents the logarithm of the relative abundance ratio in base 2. Asterisk indicates statistically significant differences (\**p* < 0.05, \*\**p* < 0.001, \*\*\**p* < 0.0001).

**S4 Fig.** Bar chart showing the top differentially abundant microbial taxa in male UTI patients and male healthy individuals. The log2-transformed relative abundance represents the logarithm of the relative abundance ratio in base 2. Asterisk indicates statistically significant differences (\**p* < 0.05, \*\**p* < 0.001, \*\*\**p* < 0.0001).

**S1 Table. Metadata of all samples included in the study**

**S2 Table. Abundance count data**

**S3 Table. Differential abundance species**

**S4 Table. Differential abundance pathway**

**S5 Table. Top differentially abundant pathway**

## Notes

### Competing Interest Statement

The authors have declared no competing interest.

## References

1. Bader MS, Loeb M, Brooks AA. An update on the management of urinary tract infections in the era of antimicrobial resistance. Postgraduate medicine. 2017 Feb 17;129(2):242–58.

2. Mancuso G, Midiri A, Gerace E, Marra M, Zummo S, Biondo C. Urinary tract infections: the current scenario and future prospects. Pathogens. 2023 Apr 20;12(4):623.

3. Flores-Mireles AL, Walker JN, Caparon M, Hultgren SJ. Urinary tract infections: epidemiology, mechanisms of infection and treatment options. Nature reviews microbiology. 2015 May;13(5):269–84.

4. Cipriani C, Carilli M, Rizzo M, Miele MT, Sinibaldi-Vallebona P, Matteucci C, et al. Bioactive compounds as alternative approaches for preventing urinary tract infections in the era of antibiotic resistance. Antibiotics. 2025 Feb 1;14(2):144.

5. Kaur R, Kaur R. Symptoms, risk factors, diagnosis and treatment of urinary tract infections. Postgraduate medical journal. 2021 Dec;97(1154):803–12.

6. Zeng Z, Zhan J, Zhang K, Chen H, Cheng S. Global, regional, and national burden of urinary tract infections from 1990 to 2019: an analysis of the global burden of disease study 2019. World Journal of Urology. 2022 Mar;40(3):755–63.

7. Zhu C, Wang DQ, Zi H, Huang Q, Gu JM, Li LY, et al. Epidemiological trends of urinary tract infections, urolithiasis and benign prostatic hyperplasia in 203 countries and territories from 1990 to 2019. Military Medical Research. 2021 Dec 9;8(1):64.

8. Saraiva A, Raheem D, Roy PR, BinMowyna MN, Romão B, Alarifi SN, et al. Probiotics and Plant-Based Foods as Preventive Agents of Urinary Tract Infection: A Narrative Review of Possible Mechanisms Related to Health. Nutrients. 2025 Mar 11;17(6):986.

9. Moustafa A, Li W, Singh H, Moncera KJ, Torralba MG, Yu Y, et al. Microbial metagenome of urinary tract infection. Scientific reports. 2018 Mar 12;8(1):4333.

10. Thomas S, Dunn CD, Campbell LJ, Strand DW, Vezina CM, Bjorling DE, et al. A multi-omic investigation of male lower urinary tract symptoms: Potential role for JC virus. PLoS One. 2021 Feb 25;16(2):e0246266.

11. Nadifah F, Artama WT, Daryono BS, Retnaningrum E. Characterization of the urogenital microbiome in patients with urinary tract infections. Indonesian Journal of Biotechnology. 2022;27(3):142–150.

12. Marshall CW, Kurs-Lasky M, McElheny CL, Bridwell S, Liu H, Shaikh N. Performance of conventional urine culture compared to 16S rRNA gene amplicon sequencing in children with suspected urinary tract infection. Microbiology Spectrum. 2021 Dec 22;9(3):e01861–21.

13. Lewis DA, Brown R, Williams J, White P, Jacobson SK, Marchesi JR, et al. The human urinary microbiome; bacterial DNA in voided urine of asymptomatic adults. Frontiers in cellular and infection microbiology. 2013 Aug 15;3:41.

14. Siddiqui H, Nederbragt AJ, Lagesen K, Jeansson SL, Jakobsen KS. Assessing diversity of the female urine microbiota by high throughput sequencing of 16S rDNA amplicons. BMC microbiology. 2011 Nov 2;11(1):244.

15. Gottschick C, Deng ZL, Vital M, Masur C, Abels C, Pieper DH, Wagner-Döbler I. The urinary microbiota of men and women and its changes in women during bacterial vaginosis and antibiotic treatment. Microbiome. 2017 Aug 14;5(1):99.16.

16. Sahil R, Pal V, Kharat AS, Jain M. A Multi-Omics Meta-Analysis of Rhizosphere Microbiome Reveals Growth-Promoting Marker Bacteria at Different Stages of Legume Development. Plant, Cell & Environment. 2025 Feb 14.

17. Ewels P, Magnusson M, Lundin S, Käller M. MultiQC: summarize analysis results for multiple tools and samples in a single report. Bioinformatics. 2016 Oct 1;32(19):3047–8.

18. Chen S, Zhou Y, Chen Y, Gu J. fastp: an ultra-fast all-in-one FASTQ preprocessor. Bioinformatics. 2018 Sep 1;34(17):i884–90.19.

19. Callahan BJ, McMurdie PJ, Rosen MJ, Han AW, Johnson AJ, Holmes SP. DADA2: High-resolution sample inference from Illumina amplicon data. Nature methods. 2016 Jul;13(7):581–3.

20. Lahti, L., and S. Shetty 2018. Microbiome R Package [WWW Document]. http://microbiome.github.io.

21. Ma S, Shungin D, Mallick H, Schirmer M, Nguyen LH, Kolde R, et al. Population structure discovery in meta-analyzed microbial communities and inflammatory bowel disease using MMUPHin. Genome biology. 2022 Oct 3;23(1):208.

22. McMurdie PJ, Holmes S. phyloseq: an R package for reproducible interactive analysis and graphics of microbiome census data. PloS one. 2013 Apr 22;8(4):e61217.

23. Dray S, Dufour AB. The ade4 package: implementing the duality diagram for ecologists. Journal of statistical software. 2007 Sep 30;22:1–20.

24. Douglas GM, Maffei VJ, Zaneveld JR, Yurgel SN, Brown JR, Taylor CM, et al. PICRUSt2 for prediction of metagenome functions. Nature biotechnology. 2020 Jun;38(6):685–8.

25. Robinson MD, McCarthy DJ, Smyth GK. edgeR: a Bioconductor package for differential expression analysis of digital gene expression data. bioinformatics. 2010 Jan 1;26(1):139–40.

26. Colella M, Topi S, Palmirotta R, D’Agostino D, Charitos IA, Lovero R, Santacroce L. An overview of the microbiota of the human urinary tract in health and disease: current issues and perspectives. Life. 2023 Jun 30;13(7):1486.

27. Kawalec A, Zwolińska D. Emerging role of microbiome in the prevention of urinary tract infections in children. International Journal of Molecular Sciences. 2022 Jan 14;23(2):870.

28. Saenz CN, Neugent ML, De Nisco NJ. The human urinary microbiome and its potential role in urinary tract infections. European Urology Focus. 2024 Dec 1;10(6):889–92.

29. Neugent ML, Hulyalkar NV, Nguyen VH, Zimmern PE, De Nisco NJ. Advances in understanding the human urinary microbiome and its potential role in urinary tract infection. MBio. 2020 Apr 28;11(2):10–128.

30. Antunes-Lopes T, Vale L, Coelho AM, Silva C, Rieken M, Geavlete B, Rashid T, Rahnama’i SM, Cornu JN, Marcelissen T. The role of urinary microbiota in lower urinary tract dysfunction: a systematic review. European urology focus. 2020 Mar 15;6(2):361–9.

31. Pan M, Hidalgo-Cantabrana C, Goh YJ, Sanozky-Dawes R, Barrangou R. Comparative analysis of Lactobacillus gasseri and Lactobacillus crispatus isolated from human urogenital and gastrointestinal tracts. Frontiers in microbiology. 2020 Jan 22;10:499288.

32. Petrova MI, Lievens E, Malik S, Imholz N, Lebeer S. Lactobacillus species as biomarkers and agents that can promote various aspects of vaginal health. Frontiers in physiology. 2015 Mar 25;6:81.

33. Bradshaw CS, Plummer EL, Muzny CA, Mitchell CM, Fredricks DN, Herbst-Kralovetz MM, Vodstrcil LA. Bacterial vaginosis. Nature Reviews Disease Primers. 2025 Jun 19;11(1):43.

34. Kim JM, Park YJ. Lactobacillus and urine microbiome in association with urinary tract infections and bacterial vaginosis. Urogenital Tract Infection. 2018 Apr 1;13(1):7–13.

